# Heterogeneity in the transcriptional response of the human pathogen *Aspergillus fumigatus* to the antifungal agent caspofungin

**DOI:** 10.1101/2021.07.15.452449

**Authors:** Ana Cristina Colabardini, Fang Wang, Zhiqiang Dong, Lakhansing Pardeshi, Marina Campos Rocha, Jonas Henrique Costa, Thaila Fernanda dos Reis, Alec Brown, Qais Z. Jaber, Micha Fridman, Taicia Fill, Antonis Rokas, Iran Malavazi, Koon Ho Wong, Gustavo Henrique Goldman

## Abstract

*Aspergillus fumigatus* is the main causative agent of invasive pulmonary aspergillosis (IPA), a severe disease that affects immunosuppressed patients worldwide. The fungistatic drug caspofungin is the second line therapy against IPA but has increasingly been used against clinical strains that are resistant to azoles, the first line antifungal therapy. In high concentrations, caspofungin induces a tolerance phenotype with partial reestablishment of fungal growth called caspofungin paradoxical effect (CPE), resulting from a change in the composition of the cell wall. An increasing number of studies has shown that different isolates of *A. fumigatus* exhibit phenotypic heterogeneity, including heterogeneity in their CPE response. To gain insights into the underlying molecular mechanisms of CPE response heterogeneity, we analyzed the transcriptomes of two *A. fumigatus* reference strains, Af293 and CEA17, exposed to low and high caspofungin concentrations. We found that there is a core transcriptional response that involves genes related to cell wall remodeling processes, mitochondrial function, transmembrane transport, and amino acid and ergosterol metabolism, and a variable response related to secondary metabolite (SM) biosynthesis and iron homeostasis. Specifically, we show here that the overexpression of a SM pathway that works as an iron chelator extinguishes the CPE in both backgrounds, whereas iron depletion is detrimental for the CPE in Af293 but not in CEA17. We next investigated the function of the transcription factor CrzA, whose deletion was previously shown to result in heterogeneity in the CPE response of the Af293 and CEA17 strains. We found that CrzA constitutively binds to and modulates the expression of several genes related to processes involved in caspofungin tolerance, and that *crzA* deletion differentially impacts the SM production and growth of Af293 and CEA17. As opposed to the Δ*crzA*^CEA17^ mutant, the Δ*crzA*^Af293^ mutant fails to activate cell wall remodeling genes upon caspofungin exposure, which most likely severely affects its macrostructure and extinguishes its CPE. This work describes how heterogeneity in the response to an antifungal agent between *A. fumigatus* strains stems from heterogeneity in the function of a transcription factor and its downstream target genes.

## Introduction

Invasive pulmonary aspergillosis (IPA) is a severe opportunistic fungal disease (Bassetti and Bouza, 2017). IPA is a major infectious complication with mortality rates as high as 90% depending on the immunological status of the host and the tissue affected (Taccone et al., 2015). Patients with acute and chronic leukemia and those that have received allogenic hematopoietic stem cell transplants are the most susceptible to IPA because neutropenia and steroid-upregulated immunosuppression are the disease’s main risk factors (Dragonetti et al., 2017). The major causative agent of IPA is the filamentous fungus *Aspergillus fumigatus* (Denning, 1998). *A. fumigatus* is a ubiquitous saprotrophic fungus that produces small hydrophobic airborne asexual spores or conidia (Abad et al., 2010). Humans inhale ∼100 to ∼1,000 *A. fumigatus* conidia every day, which can penetrate deep in bronchioles and alveolar space (Nayak et al., 2018). In healthy individuals, conidia are rapidly eliminated by mucociliary clearance and pulmonary epithelial cells (Margalit and Kavanagh, 2015). The remaining conidia are phagocytized and killed by alveolar macrophages, which initiate an inflammatory response at the infection sites. Recruited neutrophils may attach to swollen hyphae and degranulate, damaging them by oxidative and non-oxidative mechanisms (Madan et al., 1997). Pre-existing epithelial damage favors conidial attachment and germination on pulmonary epithelium; additionally, both conidia and hyphae can secrete proteases and secondary metabolites (SMs) that damage the epithelium and modulate the immune response, promoting infection establishment and evasion of host defenses in immunocompromised patients (Myers et al., 2017).

*A. fumigatus* has a cosmopolitan distribution, and its environmental and clinical isolates are genotypically and phenotypically diverse, including their pathogenic potential (Alshareef and Robson, 2014; Kowalski et al., 2016; Ries et al., 2019; Dos Santos et al., 2020; Steenwyk (a) et al., 2020). The two reference strains used in experimental studies, Af293 and CEA10 (Bertuzzi et al., 2021), which are clinical isolates, displayed significant differences in terms of physiological responses to abiotic stimuli and virulence in a murine model of IPA (reviewed by Keller, 2017). Their genomes are about 99.8% identical at the nucleotide level. Although many of the strains’ genomic differences stem from single nucleotide polymorphisms (SNPs), the two strains also differ in certain regions that harbor pseudogenes, repetitive elements, and genes that encode proteins involved in heterokaryon incompatibility (Nierman et al., 2005; Fedorova et al. 2008). The Af293 strain contains two additional biosynthetic gene clusters (BGCs) involved in SMs production (Rokas, 2007, Fedorova et al., 2008; Lind et al., 2017), while CEA10 carries multiple SNPs within 26 conserved BGCs, resulting in lower production of most of these compounds (Knox et al., 2016, Throckmorton et al., 2016). Af293 strain conidia are more resistant to antifungals exposure under *in vitro* conditions (Rokas et al., 2007) and less susceptible to the host immune system (Rosowski et al., 2018), however the Af293 strain has decreased virulence and growth in hypoxia than CEA10 (Kowalski et al., 2016; Caffrey-Carr, et al., 2017). Kowalski and collaborators (2016) showed that serial passages of Af293 at low oxygen levels increased the strain’s resistance to hypoxia and virulence, highlighting the role of adaptation to environments in *A. fumigatus* heterogeneity and virulence (Kowalski et al., 2016).

Azoles are fungicidal drugs for *A. fumigatus* and the main antifungal agents recommended for IPA (Ostrosky-Zeichner and Al-Obaidi, 2017), whereas the fungistatic echinocandins represent a second line therapy and have been used in combined therapies since azole resistant strains emerged (Mavridou et al., 2015). Caspofungin (CSP), the first echinocandin to be made available in the USA (Rybowicz and Gurk-Turner, 2002), acts by noncompetitively inhibiting the fungal β-1,3-glucan synthase (Fks1), an enzyme required for the biosynthesis of β-1,3-glucan, a primary fungal cell wall carbohydrate (Perlin, 2015). However, in a certain range of higher concentrations, CSP exhibits reduced antifungal activity, a phenomenon known as “caspofungin paradoxical effect” (CPE), known as a cellular tolerance response that alters the cell wall content and fungal growth (Aruanno et al., 2019). An overproduction of chitin stabilizes the cell wall at early stages of CSP exposure, however this increase alone does not confer resistance to the drug (Stevens, 2004). CSP induces Fks1 relocation from the hyphal tips to the vacuole, which is reversed after continuous exposure to high drug concentrations (Moreno-Velásquez et al., 2017). The drug also increases oxygen consumption and ATP production (Aruanno et al., 2019), while it generates reactive oxygen species (ROS) in the mitochondria, possibly as an off-target effect (Satish et al., 2019). Mitochondrial ROS accumulation alters the plasma membrane lipid composition (Shekova et al., 2017), probably creating a specific niche for the Fks1 enzyme, protecting it, preventing CSP from binding to its target and therefore restoring its activity (Satish et al., 2019; Valero et al., 2020).

CSP exposure increases the intracellular calcium (Ca^2+^) concentration, which binds to calmodulin and activates calcineurin by phosphorylation (Juvvadi et al., 2014). Activated calcineurin then dephosphorylates the transcription factor (TF) CrzA, which translocates to the nucleus (Ries et al., 2017) and regulates the activation of several stress responses and cell wall modifications (Soriani et al., 2008; Ries et al., 2017). Mitochondrial respiratory chain inhibition by rotenone abolishes CPE, as ATP is required for the uptake of extracellular Ca^2+^ and activation of Ca^2+^/calcineurin pathway (Aruanno et al., 2019). However, while *crzA* deletion in the clinical strain Af293 results in CPE loss (Fortwendel et al., 2010), the same does not happen in the CEA10 background (e.g., in a derivative strain called CEA17) (Ries et al., 2017), demonstrating intra-species differences or CPE heterogeneity.

In this work, we investigate the impact of low and high concentrations of CSP on transcriptional profiling of the *A. fumigatus* reference strains, Af293 and CEA17. We show that CSP exposure results in a core transcriptional response that involves genes related to cell wall remodeling mitochondrial respiratory complexes, transmembrane transport, amino acid and ergosterol metabolism. In addition, we detected differences between Af293 and CEA17 in the modulation of siderophore biosynthetic genes, and the most noticeable difference between the two strains was the increased induction of BGCs at high CSP concentration in the Af293 strain. The overexpression of the TF responsible for the SM hexadehydroasthechome (HAS) biosynthesis, which acts as an iron chelator, abolished the CPE in both backgrounds, while iron absence differentially affected Af293 and CEA17. Furthermore, to investigate the role of the TF CrzA in CPE heterogeneity, we first found that it constitutively targets several genes involved on CSP tolerance in both strains. However, deletion of *crzA* affects the two strains differently in the presence or absence of CSP. Specifically, we observed a growth defect and increased SM production in Δ*crzA*^Af293^ at standard conditions, and no activation of several cell wall-related genes upon CSP exposure, which probably affects its macrostructure and is detrimental for CPE induction. In contrast, the *crzA* deletion did not affect the CEA17 growth and the CSP exposure mildly affected the Δ*crzA*^CEA17^ mutant. Our results shed light into the heterogeneity of the response to CSP and CPE within the major fungal pathogen *A. fumigatus* and the pathways involved in the induction of tolerance to this important antifungal drug.

## Results

### Af293 and CEA17 exhibit CPE heterogeneity

To compare CSP sensitivity and CPE between Af293 and CEA17, we exposed these two strains to low and high CSP concentrations. It is known that CEA17 grows faster than Af293 (Bertuzzi et al., 2020), which was confirmed here by the larger diameter on solid medium (Figure 1a, left) and by the apparent higher hyphal density in liquid medium of CEA17 when compared to Af293 (Figure 1b, left). *A. fumigatus* conidia incubated in the presence of low CSP concentrations can germinate at slow rates as short and hyper-branched hyphae (Moreno-Velásquez et al., 2017). In solid medium, CSP achieves the maximum growth inhibition at 1 µg/ml (Figure 1a, middle), while in liquid medium this effect was observed at 0.2 µg/ml (Figure 1b, middle). These two concentrations were already defined as the CSP minimum effective concentrations (MEC) (Ries et al., 2017) for solid and liquid medium. At high CSP concentrations, long hyphae that resemble the growth in the absence of drug emerge from the slow-growing microcolonies, thus defining the paradoxical growth. This effect was observed at 8 µg/ml in solid medium (Figure 1a, right) and 2 μg/ml in liquid medium (Figure 1b, right) (Ries *et al*., 2017). Consistent with the growth differences, the CEA17 hyphae appeared denser at this concentration when compared to Af293 (Figure 1a, b). Surprisingly, the incubation at high CSP concentrations (8 µg/ml) on solid medium resulted in larger diameter of Af293 colony when compared to CEA17. At this concentration, the Af293 strain recovered ∼50% of its original growth, while the CEA17 strain recovered only ∼30% (Figure 1c), revealing a more pronounced paradoxical growth of Af293. These observations demonstrated that there is heterogeneity in CPE between these two *A. fumigatus* strains.

**Figure 1:**
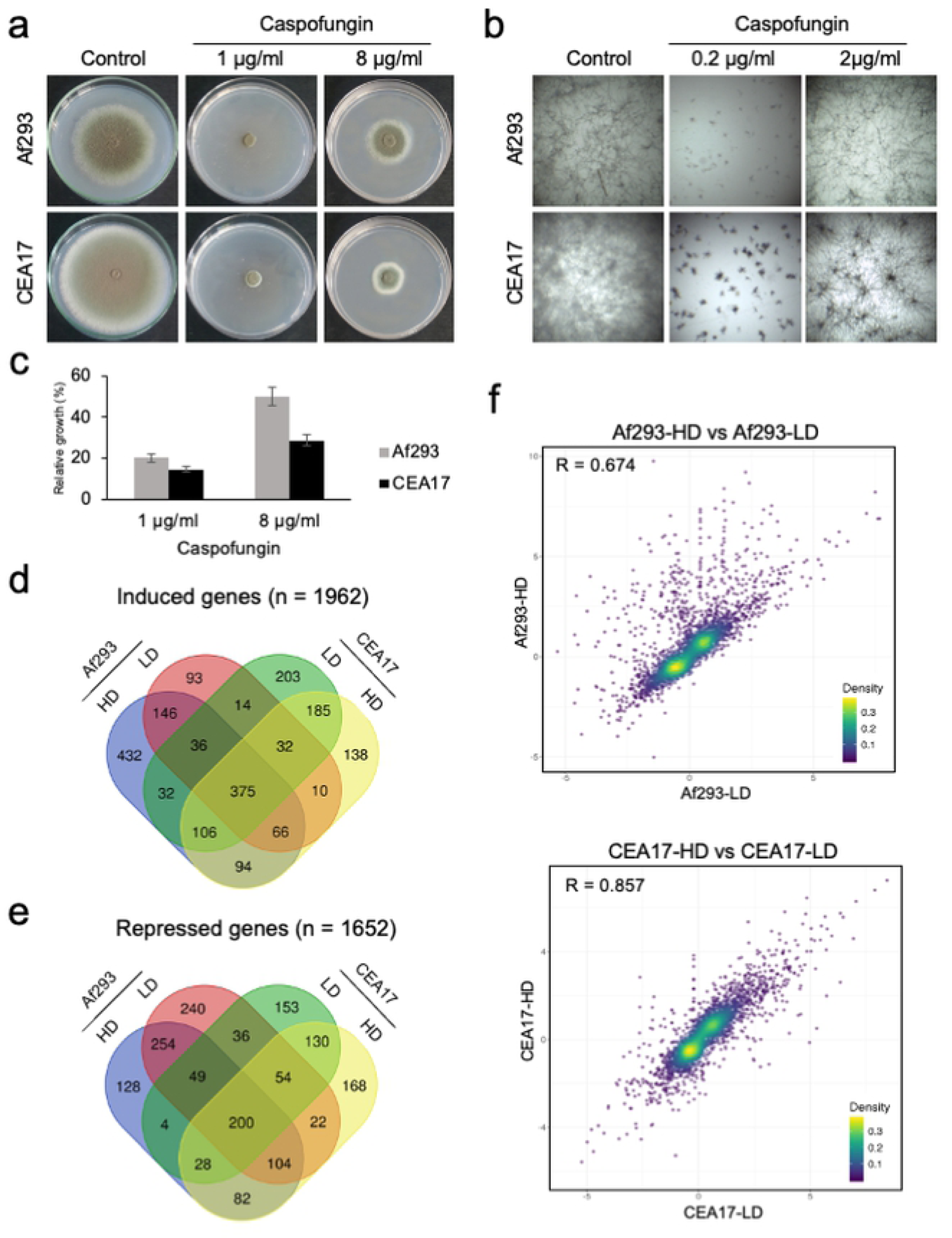
***A. fumigatus* exposed to CSP.** (a,b) Images of Af293 and CEA17 strains growth in solid (a) and liquid (b) medium during exposure to 1 and 8 µg/ml of CSP (a) and 0.2 and 2 µg/ml of CSP (b), (c) and a histogram representing the relative growth of Af293 and CEA17 when exposed to 1 or 8 µg/ml of CSP, (d-e) Venn diagrams representing the common and unique genes induced (d) and repressed (e) in Af293 and CEA17 after exposure to LD and HD of CSP, (f) scatter plots representing the correlation between the relative expression (log_2_ fold-change) of the transcriptome pairs.

### Af293 and CEA17 have dissimilar transcriptional response to CSP

To detect possible reasons for the heterogeneity observed between Af293 and CEA17, we characterized their transcriptional response to low and high concentrations of CSP at initial stages of drug exposure. Specifically, we analyzed their transcriptomes after 16 hours of growth in liquid medium and upon one hour exposure to 0.2 and 2 μg/ml of CSP, herein referred as low dose (LD) and high dose (HD), respectively. RNA sequencing (RNA-seq) analysis of Af293 and CEA17 transcriptomes (Supplementary File 1) revealed that exposure to LD and HD of CSP affected the expression of 3,614 genes in the two strains. A total of 1,962 genes were induced by LD and HD, of which 375 were commonly induced in the two strains by both doses (Figure 1d). A total of 1,652 genes were repressed by LD and HD, of which 200 were commonly repressed in the two strains by both doses (Figure 1e). Interestingly, we observed a larger set of genes commonly modulated in both strains by HD (94 induced and 82 repressed) when compared to LD (14 induced and 36 repressed) (Figure 1d-e), suggesting that the exposure to HD of CSP resulted in more changes in the transcriptional program. It is also noteworthy that we detected three times more genes (n = 432) induced exclusively in Af293 by HD when compared to in CEA17 under the same condition (n = 138) (Figure 1d), representing the first marked difference between the transcriptomes of the two strains.

To assess the correlation between the relative expression values (log_2_-fold change) of all genes from Af293 and CEA17 when exposed to LD and HD, we analyzed the Pearson’s coefficient (R) for all pairs of strains and conditions. The highest correlation value was observed between CEA17-HD and CEA17-LD (R = 0.857), followed by Af293-HD and Af293-LD (R = 0.674) (Figure 1f), suggesting that the CEA17 transcriptome is less affected with the change in dose than the Af293 transcriptome. When we compared the two strains, we observed higher correlations between Af293-LD and CEA17 [LD (R = 0.622) and HD (R = 0.665)] than between Af293-HD and CEA17 [LD (R = 0.444) and HD (R = 0.564)] (Supplementary Figure 1). Consistent with the larger set of genes induced exclusively in Af293 by HD, all Pearson’s coefficient values were lower when the comparisons were made against Af293-HD, which suggests the transcriptional response of Af293 at HD is the most different.

### CSP induces cell wall remodeling pathway genes in Af923 and CEA17 strains

In light of the above observations, we next asked which genes were commonly induced by CSP exposure in both strains. We found that induced genes were involved in cell wall organization or biogenesis, including those that encode α-glucan synthases (*ags1* and *ags3)*, ß-1,3-glucan synthase (*fks1*), 1,3-ß-glucanosyltransferases (*bgt1* and *bgt2*), chitin synthases (*chsA, chsG*), chitosanases *(csn* and *csnC),* endo-glucanases (*eng2*, *eng4, eng7* and *eng8*), exoglucanases (*exg15, exg16, exg19 and exgO*), glucanosyltransferases (*gel1, gel4, gel5 and gel7),* mannosidase (*man*), transglycolase (*crh1*), UDP-glucose 4-epimerase (*uge4*), endopeptidase (*yap2*) and xylanase (*xynf11a*) (Figure 2a and Supplementary Table 1). We also detected induction of five co-regulated genes (*gtb3, agd3, ega3, sph3, uge3*) that encode the proteins required for galactosaminogalactan (GAG) biosynthesis, one gene (*dfg1*) that is involved in the covalent binding of galactomannan (GM) to the β-(1,3)-glucan–chitin layer (Figure 2a and Supplementary Table 1), genes that encode heat shock proteins (HSPs) (e.g., *hsp90*), five genes that encode conidial hydrophobins (*rodA, rodC, rodD, rodE* and *rodF*) and several genes encoding proteins involved in calcium homeostasis (*anxc3.1, anxc3.2, anxc4, cch1, pmcA, pmcB* and *pmcC*) (Supplementary Table 1).

**Figure 2:**
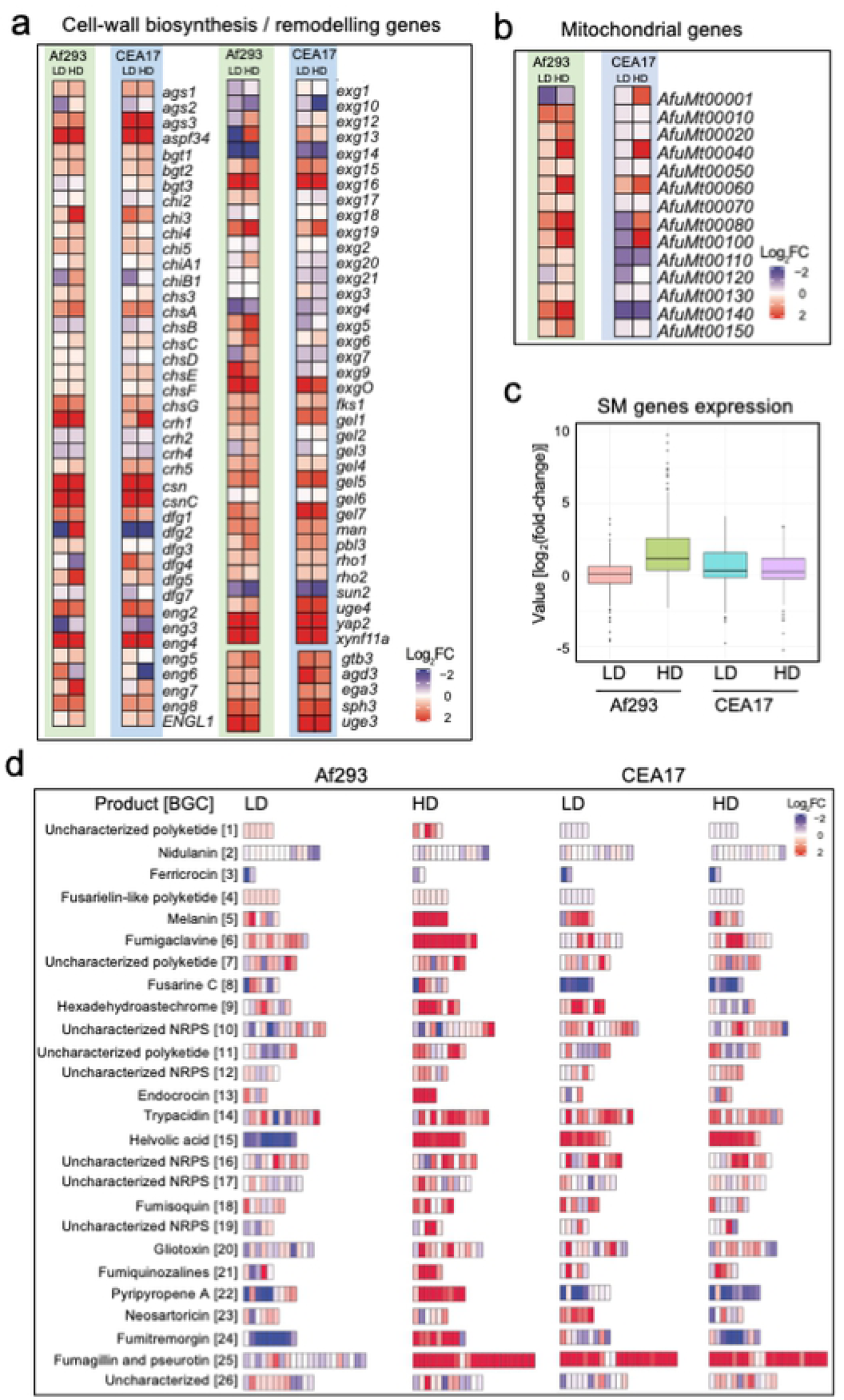
Genes induced by CSP exposure. (a,b) Heatmaps representing the relative expression of specific gene groups after exposure to LD and HD of CSP, (c) a boxplot representing the relative expression of SM genes in each transcriptome, and (e) heatmaps representing the relative expression of 26 *A. fumigatus* BGCs in Af293 and CEA17 after exposure to LD and HD of CSP.

Several cell wall remodeling enzymes are attached to the cell membrane via glycosylphosphatidylinositol (GPI) anchors, whose biosynthesis requires mevalonate. Here, we detected induction of the *hmg2* gene, which encodes a hydroxymethylglutaryl-CoA (HMG-CoA) reductase involved in mevalonate biosynthesis (Supplementary Table 1). In eukaryotic cells, the endoplasmic reticulum (ER) is responsible for the synthesis of membrane proteins, including GPIs (Lopez et al., 2019). When the burden of nascent unfolded and misfolded proteins in the ER increases beyond its processing capacity, an unfolded protein response (UPR) is activated (Ron and Walter, 2007). Given the involvement of GPIs and the ER, caspofungin-induced processes are expected to induce the UPR. Accordingly, we detected induction of genes that encode proteins involved in the UPR, such as *bipA, casA, casB, clxA, hacA* and *ireA* (Supplementary Table 1). In general, the genes described in this section were similarly induced by LD and HD in the two strains and can be thought as representing the core CSP response in *A. fumigatus*.

### The mitochondrial respiratory chain biosynthetic genes are highly induced at high dose of CSP

It was previously shown that CSP induces the expression of the genes of the mitochondrial respiratory chain (MRC) in CEA17 and that mitochondrial activity is essential for the CPE (Cagas et al., 2011; Conrad et al., 2018; Aruanno et al., 2019; Valero et al., 2020). Here, we detected the four genes encoding the components of the mitochondrial respiratory chain (MRC) complexes IV (AfuMt00040, AfuMt00060 and AfuMt00080) and V (AfuMt00100) were highly induced by HD in Af293 and CEA17 strains (Figure 2b and Supplementary Table 1). Furthermore, AfuMt00010, AfuMt00020 and AfuMt00150, which encode proteins of complex I, were induced by HD in Af293, while AfuMt00001, which encodes a protein of complex III, was induced by HD only in CEA17. Notably, the induction levels were higher in Af293 treated with HD when compared to CEA17 (Figure 2b and Supplementary Table 1). Of note, three of these genes (AfuMt00010, AfuMt00080 and AfuMt00140) were induced at lower levels by LD in Af293 and one gene (AfuMt00060) was induced also at lower levels in CEA17 by LD (Figure 2b and Supplementary Table 1). These findings reinforce the relevance of mitochondria in the CPE (Cagas et al., 2011; Conrad et al., 2018; Aruanno et al., 2019; Valero et al., 2020) and suggest an explanatory mechanism for the observed differences in the growth between LD and HD, namely that a certain level of induction of MRC genes (and presumably MRC activation) must be achieved for CPE induction (since MRC genes induction were proportional to CSP concentration).

### Secondary metabolite biosynthetic genes were highly induced by HD of CSP compared to LD in the Af293 strain

After evaluating the commonly induced genes, we turned to genes that were differentially expressed in each strain. It was discussed before that HD induced a larger set of genes in Af293 (Figure 1d), which presumably contributed to the lower correlation between Af293-HD and the other conditions (Figure 1f and Supplementary Figure 1). GO enrichment analysis showed 65 out of the 432 genes induced exclusively by HD in Af293 were related to secondary metabolic processes (P-value = 1.01E-20). The analysis of the secondary metabolite (SM) biosynthetic genes expression revealed that they were more induced when compared to Af293-LD and CEA17 (LD and HD) (Figure 2c). In fact, several SM genes such as *abr1*, *abr2, easG*, *easD*, *encB*, *fmqC*, *fmqD* were induced more than 7-fold in Af293-HD, representing the highest induced genes in the whole transcriptome analysis (Supplementary Table 2). It is noteworthy that *brlA*, a gene that encodes the TF associated with conidiation and SM production (Coyle et al., 2007; Lind et al., 2018), was induced only in Af293-HD (Supplementary Table 1).

It is known that the genes encoding enzymes for the SM biosynthesis are contiguously arranged in clusters (Rokas et al., 2018; Pfannenstiel and Keller, 2019) and that *A. fumigatus* possess at least 26 SM biosynthetic gene clusters (BGCs) (Bignell et al., 2016; Lind et al., 2017). We performed a systematic analysis to identify BGCs with at least half of their genes differentially expressed. Of the 26 BGCs described by Bignell and collaborators (2016), 10 BGCs (5, 6, 9, 13, 14, 15, 21, 22, 24 and 25) were induced by HD in Af293, and 5 BGCs (5, 14, 15, 23 and 25) and 2 BGCs (15 and 25) were induced in CEA17 by LD and HD, respectively (Figure 2d and Supplementary Table 2). On the other hand, 3 BGCs (15, 22 and 24) were repressed by LD in Af293, and 2 BGCs (8 and 22) and 3 BGCs (8, 22 and 24) were repressed in CEA17 by LD and HD, respectively (Figure 2d, Supplementary Table 2). Consistent with previous work (Conrad et al., 2018), the BGC25, for the biosynthesis of fumagillin and pseurotin, was induced in CEA17 by LD and HD and in Af293 by HD. Additionally, the BGC9, for the biosynthesis of hexadehydroastechrome (HAS), was induced by HD in Af293, while the BGC8, for the biosynthesis of siderophores, was repressed by LD and HD in CEA17, and the BGC24, for the biosynthesis of fumitremorgin, was induced by HD in Af293 and repressed by LD in Af293 and by HD in CEA17 (Figure 2d and Supplementary Table 2). These results describe a striking difference not only between LD and HD exposures in Af293, but also between the Af293 and CEA17 strains, and places the differential induction of BGCs by HD as the most remarkable transcriptome difference between the two strains.

### Iron absence is detrimental for CPE in Af293

Given the BGC25 induction in Af293 and CEA17 strains upon CSP exposure, we asked if fumagillin and / or pseurotin could have a role in the CPE and / or CSP sensitivity. We tested in low (1µg/ml) and high (8µg/ml) CSP concentrations the mutants Δ*fapR*^CEA17^ and Δ*fmaA*^CEA17^, that fail to produce fumagillin/pseurotin and pseurotin, respectively, but our results showed that the deletions had no effect on CSP sensitivity or CPE (Supplementary Figure 2). As HAS has a global influence on SM genes (Wiemann et al., 2014), we then asked whether the HAS absence or overproduction influenced CSP sensitivity or CPE. Similar to the Δ*fapR*^CEA17^ and Δ*fmaA*^CEA17^ mutants, the *hasA* deletion (Δ*hasA*^CEA17^) had no effect on CEA17 growth during incubation with low and high concentrations of CSP (Supplementary Figure 2), however *hasA* overexpression (*hasA*OE) abolished the CPE in both Af293 and CEA17 strains (Figure 3a). HAS acts as an iron chelator (Li et al., 2013), so we hypothesized that iron depletion would affect the CPE in Af293 and CEA17. Surprisingly, our results showed that iron is essential for CPE in Af293, but not in CEA17 (Figure 3b).

**Figure 3:**
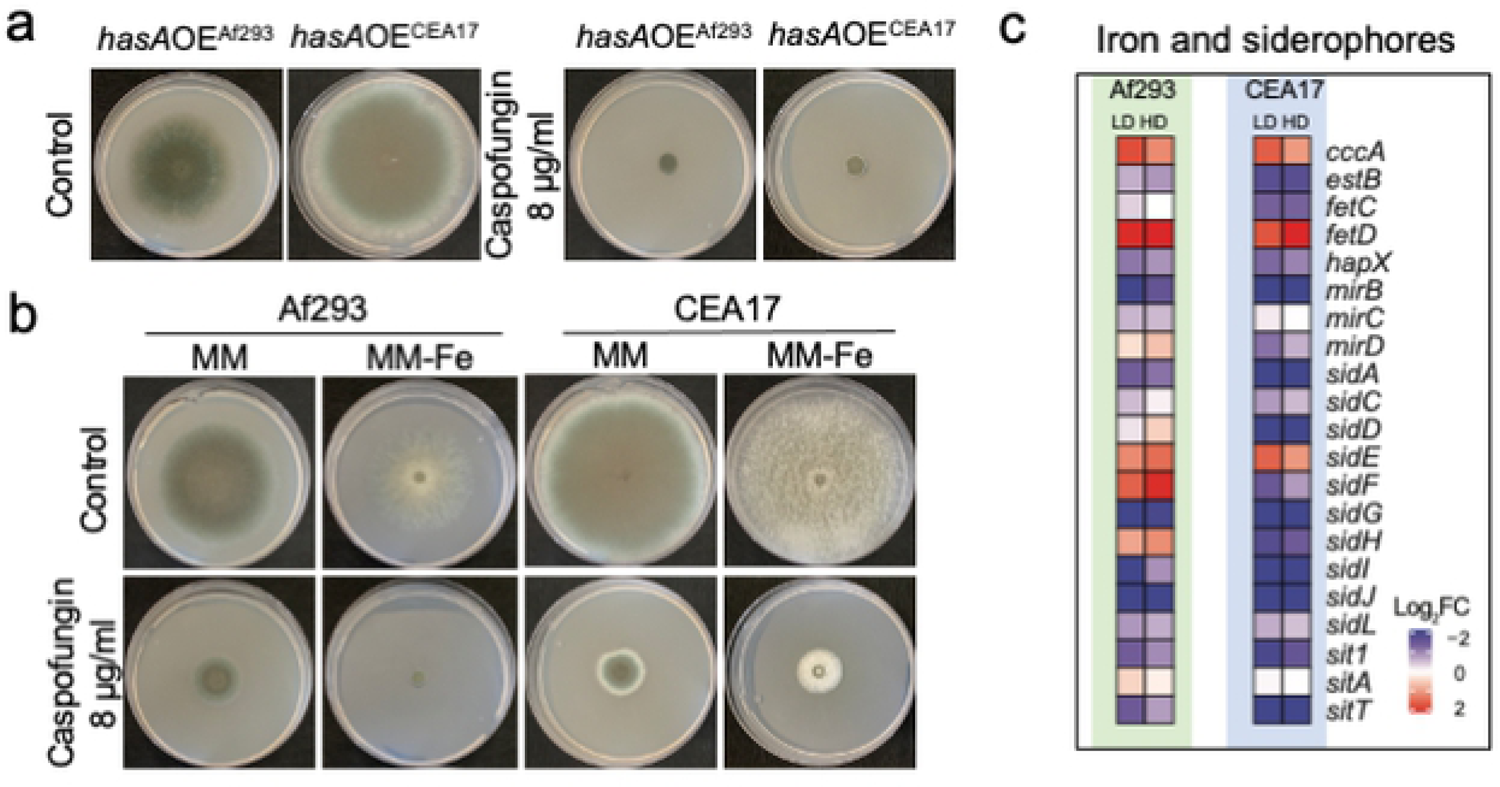
Disruption in iron homeostasis affects the CPE. (a) Images of Af293 and CEA17 strains overexpressing *hasA* grown in solid medium added with 1 and 8 µg/ml of CSP, (b) Images of Af293 and CEA17 strains grown in solid medium without iron added with 1 and 8 µg/ml of CSP, (c) heatmaps representing the relative expression of genes involved in iron homeostasis and siderophores metabolism in Af293 and CEA17 after exposure to LD and HD od CSP.

Upon iron starvation, *A. fumigatus* employs siderophore-mediated iron uptake, reductive iron assimilation, and low affinity iron uptake mechanisms to restore its iron homeostasis (Haas, 2014). Here, we detected that *sidE*, a gene essential for the biosynthesis of the siderophore fusarinine C (FsC) (Steichen et al., 2013), *cccA*, which encodes an iron transporter of the vacuolar membrane (Gsaller et al., 2012), and *fetD*, a gene normally induced during iron starvation (Yasmin et al., 2009) were induced by LD and HD in both strains. In contrast, *sidF* and *sidH*, required for the biosynthesis of the siderophore triacetylfusarinine C (TAFC) (Haas, 2014), were induced by LD and HD in Af293 and repressed in CEA17 (Figure 3c and Supplementary Table 1). Although several genes involved in the cellular response to iron starvation (*hapX* and *fetC*) and siderophore biosynthesis (*sidA, sidC, sidD, sidG, sidI, sidJ* and *sidL*) and transport (*estB, mirB, sit1* and *sitT*) were repressed in both strains Af293 and CEA17 (Figure 3c and Supplementary Table 1). These data show a conserved and a variable transcriptional response to iron starvation in *A. fumigatus* as a result of CSP exposure, indicating a difference in the iron homeostasis regulation between the two strains in response to CSP. Taken together, these data place iron metabolism as another significant difference in transcriptional regulation between the Af293 and CEA17 strains after CSP exposure.

### LD and HD of CSP commonly represses processes related to the cell membrane and translation

In our transcriptomic analysis, we detected 200 genes commonly repressed by LD and HD in both strains (Figure 1e), representing the core of the transcriptional repression by CSP. GO analysis of these genes showed that they were related to transmembrane transport, cellular amino acid metabolic process, small molecule metabolic process, plasma membrane organization and lipid metabolic process (Figure 4a). A closer inspection of the genes assigned to the later three groups revealed that several are transmembrane transporters or belong to the ergosterol biosynthetic pathway (Figure 4b and Supplementary Table 1), suggesting a possible change in membrane composition in response to CSP exposure at both concentrations. It is also noteworthy that among the genes repressed by HD in both strains (Figure 1e), we detected a conserved response that involves genes related to ribosome biogenesis, ribonucleoprotein complex assembly and tRNA metabolic process (Figure 4c and Supplementary Table 1), possibly because of the diversion of precursors from primary to secondary metabolism.

**Figure 4:**
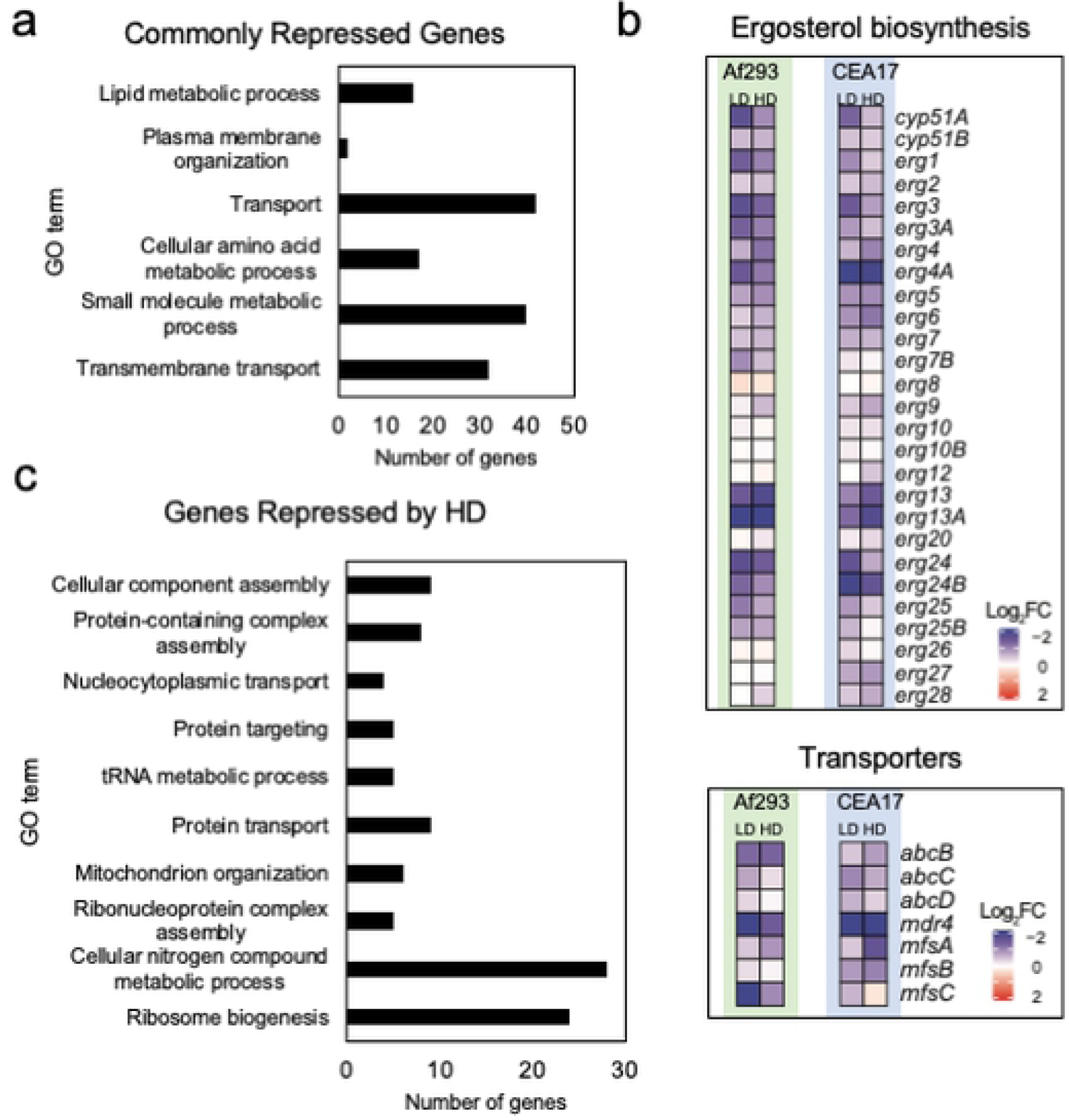
Genes repressed by CSP exposure. (a) Histogram representing the GO terms enriched among the genes commonly repressed by LD and HD of CSP in Af293 and CEA17 strains, (b) heatmaps representing the relative expression of genes involved in ergosterol metabolism and transmembrane transport after exposure to LD and HD of CSP in Af293 and CEA17 strains, (c) histogram representing the GO terms enriched among the genes commonly repressed by HD of CSP in Af293 and CEA17 strains.

### CrzA directly binds genes involved in caspofungin response in CEA17, and presumably in Af293

CSP exposure induces a spike in the concentration of cytosolic Ca^2+^ (Juvvadi et al., 2015), which results in the activation of CrzA (Ries et al., 2017), a TF involved in stress responses and cell wall remodeling (Soriani et al., 2008). In fact, CrzA participates in cell wall biosynthesis-related processes in the CEA17 strain, even in the absence of caspofungin (Ries et al., 2017). To gain an overview of the genes directly regulated by CrzA in *A. fumigatus*, the CrzA binding sites were determined genome-wide by ChIP-seq (chromatin immunoprecipitation coupled to DNA sequencing) of a CrzA-GFP tagged strain. The analysis below was performed solely in CEA17 background due to the failure to construct a CrzA-tagged Af293 strain. We detected a total of 600 genes bound by CrzA at their promoters before and after exposure to HD of CSP. Although we observed a stronger CrzA binding signal after CSP exposure at several targets (Figure 5a), all the targets were constitutively bound at the same sites (Supplementary Table 3). Enrichment analysis of the genes bound by CrzA revealed that it engages in processes related to cell wall organization or biogenesis, response to stimulus, cell communication, response to stress, response to endoplasmic reticulum stress (*i.e.* UPR), iron import into cell and siderophore metabolic process (Figure 5b). Notably, two clusters containing 7 and 5 genes, responsible for siderophore biosynthesis and transport and cell wall galactosaminogalactan (GAG) biosynthesis, respectively, were strongly bound by CrzA before and after CSP exposure (Figure 5c), while genes related to β-1,3-glucan and chitin biosynthesis (*fks1* and *chsA*), UPR, such as (*hacA*) and calcium homeostasis (*pmcA*) were more strongly bound by CrzA after CSP exposure (Figure 5d). Interestingly, we also detected a stronger CrzA binding to *crzA* promoter after CSP exposure (Figure 5d), suggesting self-regulation in this condition. Furthermore, the gene *anxc3.1* (Afu8g06700), encoding one annexin that was previously shown to implicate in *A. fumigatus* adaptation to high concentrations of calcium in a CrzA-dependent manner (Soriani et al., 2008), was also detected here as CrzA target (figure 5d). These results demonstrate that CrzA binds to promoters of genes that we showed here to be relevant for CSP response. In fact, 232 out of the 600 CrzA targets (∼38%) were modulated by CSP exposure in the CEA17 transcriptome analysis (Figure 6e), suggesting that CrzA plays relevant roles during normal growth and in response to CSP exposure.

**Figure 5:**
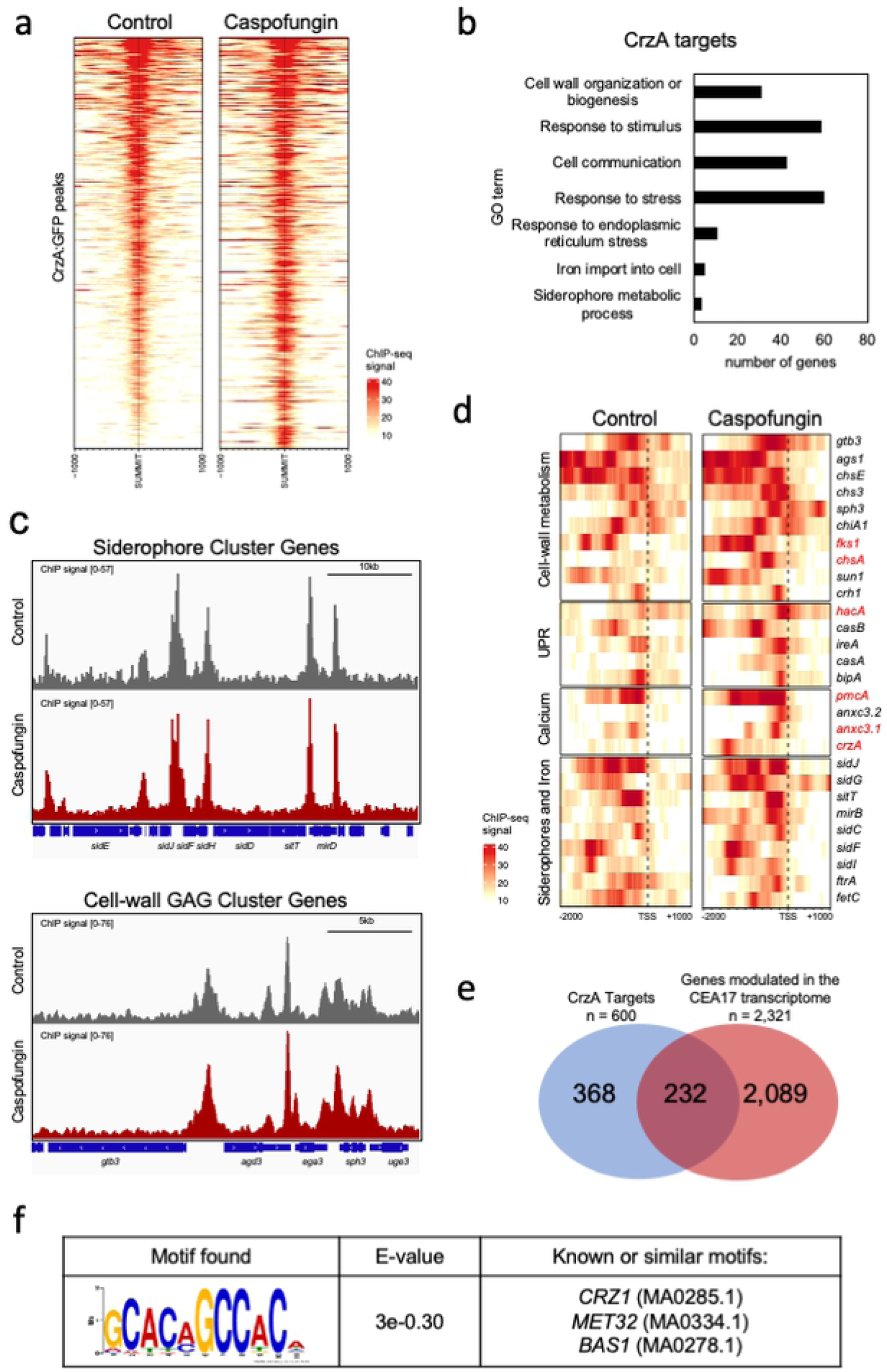
CrzA targets in CEA17. (a) Heat maps representing the intensity of CrzA binding at its targets in CEA17 before and after CSP exposure, the regions represent the peaks summit ± 1000 bp, (b) a histogram representing the GO enriched terms among the CrzA targets, (c) screenshots of the genome browser representing the peaks of CrzA binding at the promoter regions of the siderophore and GAG clusters of genes, (d) heatmaps representing the CrzA:GFP binding regions against selected genes before and after CSP exposure (d) Venn diagram representing the number of CrzA targets that overlap with the genes modulated in our transcriptome analysis, (e) motif enrichment analysis of CrzA targets.

**Figure 6:**
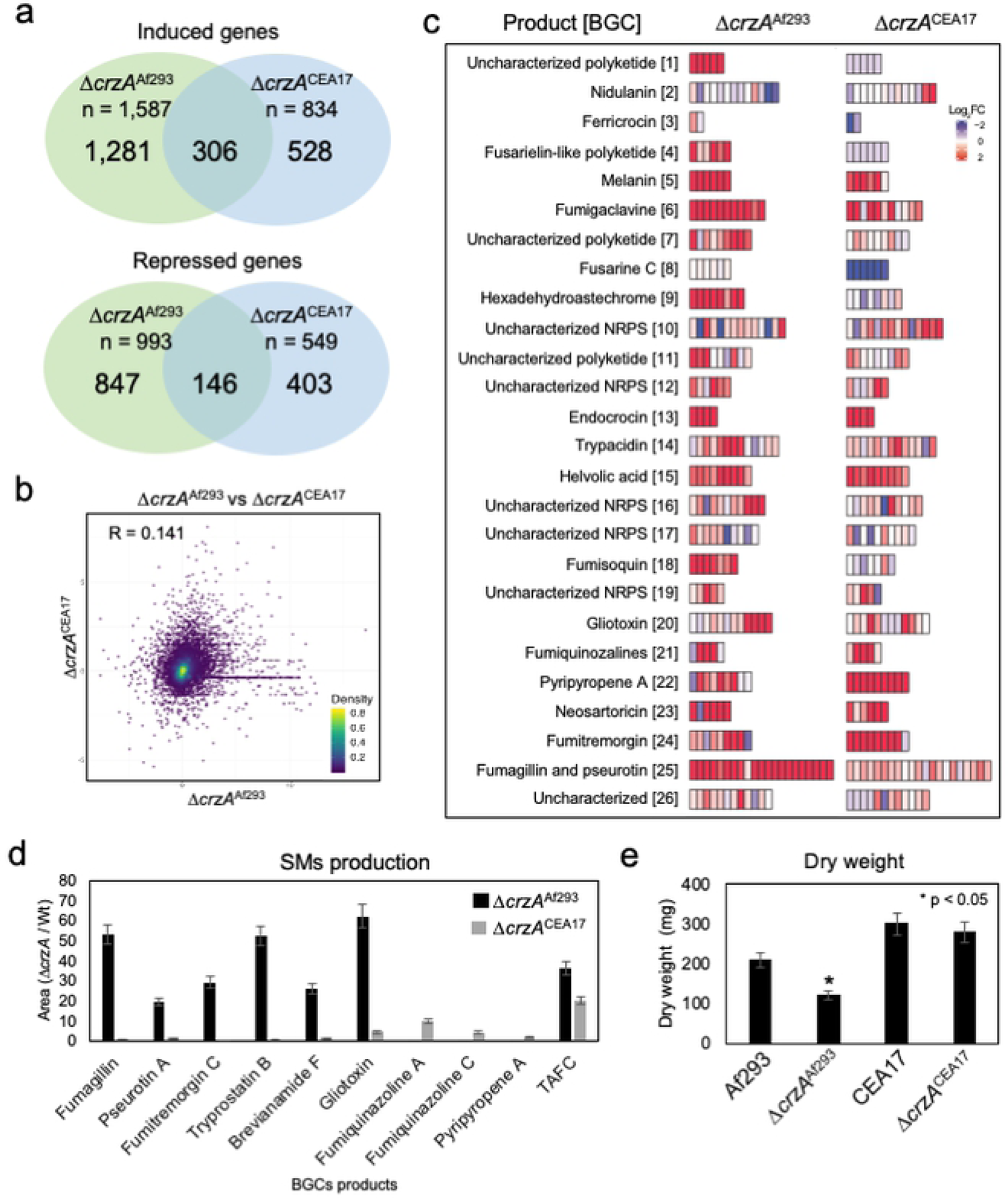
The impact of *crzA* gene deletion in Af293 and CEA17 strains. (a) Venn diagrams representing the number of genes commonly and uniquely affected by *crzA* deletion in Af293 and CEA17 strains, (b) a scatterplot representing the correlation between the relative expression of the genes as in (a), (c) heatmaps representing 26 BGCs relative expression in the *crzA* mutant strains, (d) a histogram representing the relative SM production in Δ*crzA*^Af293^ and Δ*crzA*^CEA17^, (e) a histogram representing the dry weight of the Af293, CEA17 and the Δ*crzA* mutant strains grown in standard conditions.

To determine the CrzA DNA binding motif, we performed a MEME (Multiple EM for Motif Elicitation)-ChIP analysis to search the 200 and 500 bp region surrounding the peaks identified in our ChIP-seq data (Machanick and Bailey, 2011). We observed that CrzA binds to GC-rich regions [GCC(A/T)C] (Figure 5e), a pattern similar to the *CRZ1* binding motif of *Saccharomyces cerevisiae* [5′-GNGGC(G/T)CA-3′] (Yoshimoto et al., 2002) and *Candida albicans* [5′-GGAGGC(G/A)C(T/A)G-3′] (Xu et al., 2020). Considering the differences observed between Af293 and CEA17 upon caspofungin exposure, we asked whether CrzA binding regions are similar in these strains. Based on the motif discovered, we found that 111 promoter regions contained the CrzA binding motif among the 600 targets in CEA17 (Supplementary Table 3). A comparison of these 111 promoters did not identify regions that differed between the two strains (*i.e.*, contained a different number of CrzA binding site or the CrzA binding site was in a different location within the promoter in Af293 and CEA17). Although the presence of the motif does not necessarily imply binding by CrzA, the similarity between the two strains regarding the presence of the binding motif indicates that CrzA binds in similar targets in Af293 and CEA17.

### Deletion of *crzA* slows the growth of the Af293 strain and alters secondary metabolite regulation and production

To further understand which processes are affected by CrzA and whether they are conserved between Af293 and CEA17, we analyzed the transcriptional differences on the effects of *crzA* deletion in the two strains. RNA-seq analysis of the Δ*crzA* mutants comparing to their respective parental strains revealed the induction of 1,587 genes in Δ*crzA*^Af293^ and 834 genes in Δ*crzA*^CEA17^, of which 306 were common to both strains. In addition, the repression of 993 genes in Δ*crzA*^Af293^ and 549 genes in Δ*crzA*^CEA17^ was observed of which only 146 were common to both strains (Figure 6a). Consistent with the small number of genes commonly modulated in both mutants, the correlation between the relative expression (log_2_ fold-change) of genes in the two transcriptomes was also very low (R = 0.141) (Figure 6b). GO enrichment analysis detected that 138 and 71 of the genes induced in Δ*crzA*^Af293^ and Δ*crzA*^CEA17^, respectively, were related to secondary metabolic process (Supplementary Table 4). A closer inspection of these genes showed that the BGCs involved in the biosynthesis of melanin, endocrocin, helvolic acid, fumiquinazolines and neosartoricin (Figure 6c and Supplementary Table 2) were induced in both mutants. Additionally, the BGCs involved in the biosynthesis of HAS and fumagillin and pseurotin were induced in Δ*crzA*^Af293^ and the BGCs involved in the biosynthesis of pyripyropene A and fumitremorgin were induced in Δ*crzA*^CEA17^ (Figure 6c and Supplementary Table 2).

To verify if the observed BGC induction represented increased SM production, we used LC/MS to search for different compounds in the supernatants of Δ*crzA*^Af293^ and Δ*crzA*^CEA17^ mutants and their respective parental strains. In this case, the strains were grown in minimal medium for 96 hours, the condition in which we detect the greatest SM diversity. Consistent with the transcriptome results, we detected in Δ*crzA*^Af293^ supernatants 20-50 times higher relative levels of fumagillin and pseurotin A, while fumiquinazolines A and C and pyripyropene A were detected in the Δ*crzA*^CEA17^ supernatants (Figure 6d). Interestingly, fumitremorgin C and its intermediates brevianamide F and tryprostatin B, and gliotoxin were detected at much higher levels in Δ*crzA*^Af293^ supernatants (Figure 6d), which suggests that these BGCs were induced later than 16 hours, therefore not detected in our RNA-seq. Additionally, the relative levels of the siderophore TAFC were 20-40 times increased in the supernatants of Δ*crzA*^Af293^ and Δ*crzA*^CEA17^ after 96 hours (Figure 6d), suggesting that both mutants consume more iron than their respective parental strains, most likely due to the increased SM production.

The GO analysis also revealed repression of processes related to translation, ribosome biogenesis and ribonucleoprotein complex assembly only in the Δ*crzA*^Af293^ mutant (Supplementary Table 4). Consistently, a moderate growth defect was observed in Δ*crzA*^Af293^ (Figure 6e). Taken together, these results revealed significant differences in SM production and growth when *crzA* is deleted on Af293 and CEA17 strains, suggesting that CrzA function differs between these two strains.

### Fluorescent caspofungin accumulates in the interface cell membrane/cell wall and is internalized into *A. fumigatus* vacuoles

The *crzA* deletion mutant extinguishes the CPE in Af923, but not in CEA17 (Fortwendel et al., 2010, Ries et al., 2017), so we hypothesized that CSP could be internalized at different levels depending on the background strain. Recently, Jaber and collaborators (2020) developed functional fluorescently labeled probes of CSP (FCSP). We tested one of these probes and observed that in CEA17 germlings exposed to 0.125 µg/m of FCSP for 5 min, the drug accumulates in the interface cell wall/cell membrane (Figure 7a) but after 15- and 30-min exposure, besides its presence at the cell membrane/cell wall interface, FCSP is progressively internalized into small vesicles and structures that resemble vacuoles (Figure 7a). The vacuoles were confirmed by co-staining with the blue CMAC dye (7-amino-4-chloromethylcoumarin) that enters into the fungal cytoplasm because of its hydrophobicity, and it is modified to glutathione by glutathione Stransferase (Shoji et al., 2006). In fungi, CMAC accumulates in vacuoles, most likely through the transport of glutathione transporters present on the vacuolar membrane (Shoji *et al*., 2006). The Af293, the Δ*crzA*^CEA17^, and Δ*crzA*^Af293^ strains also showed the same behavior (Supplementary Figure 3). Taken together, our results suggest a possible removal of the FCSP-Fks1 complexes by *A. fumigatus* endocytic machinery followed by a trafficking to the vacuoles. These phenotypes are similar in both strains, and they are not affected by the lack of CrzA.

**Figure 7:**
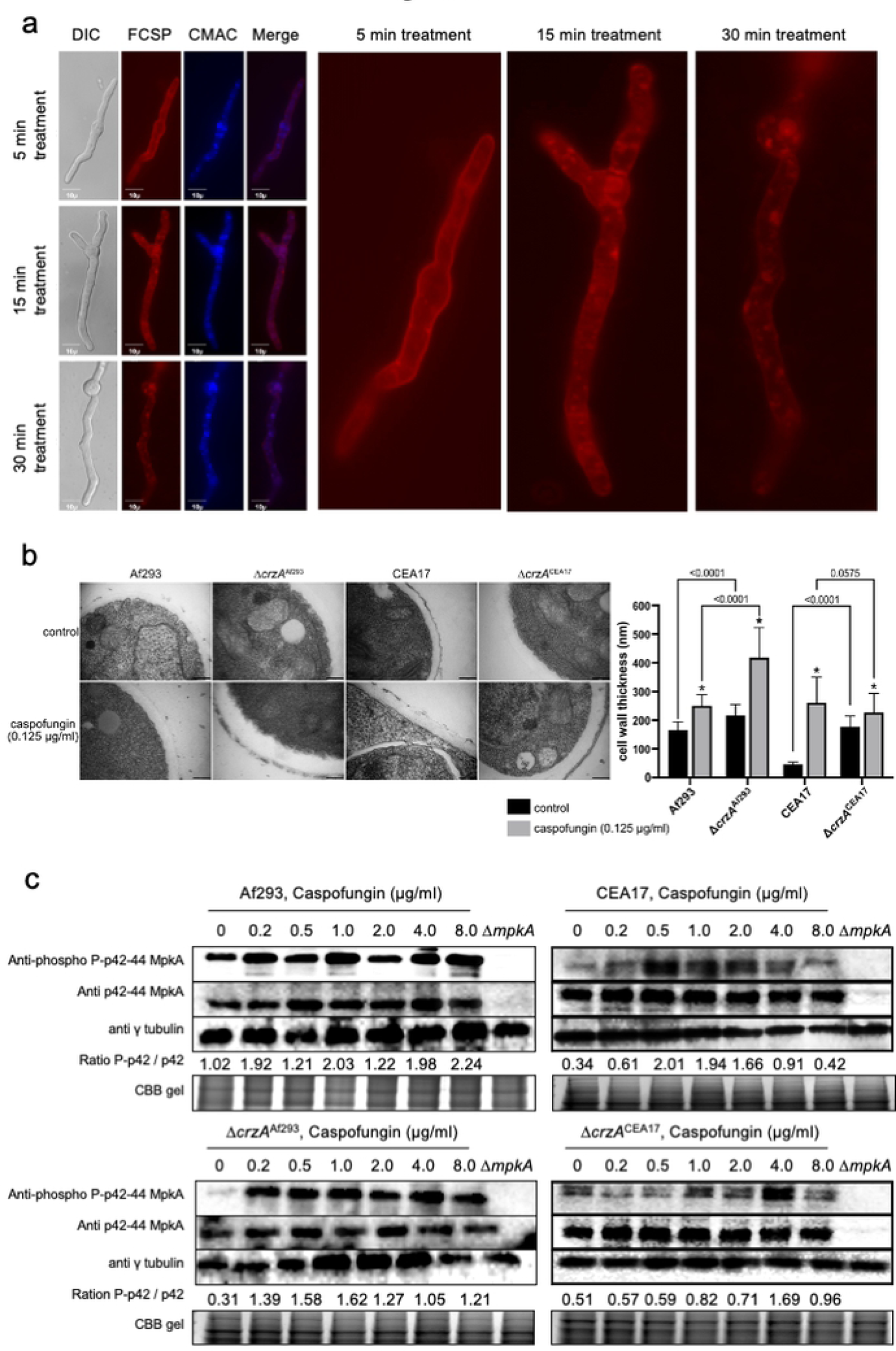
Af293, CEA17 and the Δ*crzA* mutant strains before and after CSP exposure. (a) Cellular localization determined by microscopy of the fluorescent CSP (FCSP) in the CEA17 strain after 5, 10 and 15 minutes of exposure to 0.125 µg/ml, the vacuoles were labeled with blue CMAC dye (7-amino-4-chloromethylcoumarin), (b) Transmission electron microscopy (TEM) of mycelial sections of Af293, Δ*crzA*^Af293^ CEA17, Δ*crzA*^CEA17^, strains mycelia treated or not with 0.125 µg/ml of CSP for 2 h in AMM at 37°C. Magnification bars: 200 nm. The cell wall thickness of 100 sections from different germlings (average of 4 sections per germling) was measured and plotted on the graph. Average ± SD are shown. Statistics were determined by two-way ANOVA with a Sidak’s multiple comparison test. Asterisks indicate significant differences between treated and untreated samples for the same strain (black versus gray bars, p<0.0001). All other comparisons are indicated by the bars, (c) Immunoblot assay of MpkA phosphorylation exposed to increasing CSP concentrations using anti-p44/42 MpkA or anti-44/42 MpkA antibodies to detect the phosphorylated and total MpkA, respectively. The anti-γ-tubulin antibody and a Coomassie brilliant blue (CBB) gel were used as loading controls and the signal intensities were quantified using the ImageJ software.

### Af293 and CEA17 strains have dissimilar cell walls which are differently affected by *crzA* deletion and by CSP exposure

As CSP is similarly internalized by the strains, we then considered that Δ*crzA*^Af293^ fails to activate key elements responsible for the CPE. To understand which elements are those, we analyzed the Δ*crzA*^Af293^ and Δ*crzA*^CEA17^ transcriptomes after CSP exposure (LD and HD) and compared them to the untreated ones. Of the genes shown to be important for CSP tolerance in Af293 and CEA17 (Supplementary Table 1), several genes involved in cell wall metabolism, such as *ags1, ags3, chsG, eng2, exgO, fks1, gel1* and *gel7*, were induced by caspofungin in Δ*crzA*^CEA17^, but not in Δ*crzA*^Af293^ (Supplementary Figure 4), suggesting that the Δ*crzA*^Af293^ mutant fails to activate a proper cell wall remodeling program upon CSP exposure.

To verify whether the transcriptional profile reflected any macrostructural changes in Δ*crzA*^Af293^, we analyzed by transmission electron microscopy (TEM) the cell wall of Af293 and CEA17 strains along with the Δ*crzA*^Af293^ and Δ*crzA*^CEA17^ mutants before and after exposure to a lower dose of CSP (0.125 µg/ml) (Figure 7b). In this experiment, the CSP concentration was lower than LD because the cell type used for TEM (*i.e.* germlings) is much more sensitive to the drug than the cell type used for the RNA-seq (*i.e.* mycelia). Before CSP addition, the Af293 cell wall was more than three times thicker than that of the CEA17 strain (Figure 7b). After 1 hour of CSP exposure, the Af293 and CEA17 cell walls showed similar thickness, meaning that CEA17 showed much higher increase by fold than Af293 when compared to the control not exposed to the drug (Figure 7b). Surprisingly, the Δ*crzA*^Af293^ cell wall had similar thickness than that of the Af293 strain, whereas the Δ*crzA*^CEA17^ cell wall was much thicker than that of the CEA17 strain (Figure 7b). In a comparison among all the strains, the Δ*crzA*^Af293^ cells had the thickest cell wall after CSP exposure (Figure 7b), indicating that this mutant has the cell wall most affected by CSP.

### The cell wall integrity pathway is heterogeneously activated by CSP and is partially dependent on CrzA

The cell wall integrity pathway (CWIP) is the primary signaling cascade that controls the cell wall homeostasis in fungi (Levin, 2011). In this pathway, an upstream kinase activates a three-component mitogen-activated protein kinases (MAPKs) culminating in the activation of MpkA by phosphorylation (Rocha et al., 2015). MpkA, in turn, controls the activity of the TF RlmA, which regulates the expression of genes involved in cell wall metabolism (Rocha et al., 2016; Rocha et al., 2020). As we detected heterogeneity among the Af293, CEA17, and Δ*crzA* mutant strains regarding the cell wall thickness before and after CSP exposure, we asked whether there were differences in the CWIP activation between the wild-type and the mutant strains. In the first place, we measured the MpkA phosphorylated/unphosphorylated ratio in Af293 and CEA17 strains exposed to CSP concentrations ranging from 0.2 to 8 μg/ml (Figure 7c). In Af293, we detected MpkA phosphorylation at all CSP concentrations reaching the highest levels at 8 μg/ml, while the highest level of the phosphorylation in CEA17 was observed at a lower CSP concentration (0.5 μg/ml) (Figure 7c). Furthermore, the detection of phosphorylated MpkA decreased with increasing drug concentration in CEA17 (Figure 7c), demonstrating heterogeneity in MpkA activation upon CSP exposure. Next, we verified a possible interaction between *crzA* and MpkA CWIP activation by CSP and found that the MpkA phosphorylation ratio decreased overall in the Δ*crzA*^Af293^ and Δ*crzA*^CEA17^ mutant compared to their respective parental strain, except for 0.5 µg/ml of CSP in the Δ*crzA*^Af293^ mutant and for 4.0 µg/ml of CSP in the Δ*crzA*^CEA17^ mutant. These results suggest that, although not essential, *crzA* is important for the MpkA CWIP activation in *A. fumigatus*.

## Discussion

Fungal cell walls have evolved numerous mechanisms to protect cell integrity. Although most of their building blocks are conserved, there is variation in their amounts and in the way they are organized, resulting in high diversity. Since cell walls are absent in humans, antifungals that target the production of their components, such as CSP, are more selective and less toxic when compared to drugs that target the fungal membrane. Several recent studies have uncovered variation in virulence and physiological responses to several stimuli between the two *A. fumigatus* model strains, Af293 and CEA17 (Keller, 2017; Kowalski et al., 2016; Fuller et al., 2016; Steenwyk et al., 2020a; Steenwyk et al., 2020b; Dos Santos et al., 2020; Ries et al., 2019). Investigation and comparison of the molecular mechanisms involved in this heterogeneity can reveal mechanisms (and diversity in mechanisms) to cope with different kinds of stress. Here we demonstrate that the exposure to low concentrations of CSP inhibits Af293 and CEA17 growth and makes their cell walls equally thicker. At higher concentrations, both strains had their growth partially recovered so that Af293 had a more pronounced CPE compared to CEA17, most likely due to the differences in the cell wall composition and transcriptional program. Even though both strains had a larger number of genes induced at higher concentration of drug, exposure to low and high CSP concentrations resulted in a core response that involved the induction of several genes involved in cell wall assembly and remodeling. Although this transcriptional response has already been reported (Altwasser et al., 2015; Conrad et al., 2018), our study shows that this response is conserved in the two strains and occurs in a range of CSP concentrations, implying that this compensatory mechanism alone is not the main CPE effector.

Most of the cell wall synthases and remodeling enzymes are attached to the membranes via GPI-anchors, whose biosynthesis occurs in the ER (Li et al., 2018), indicating that the activation of cell wall remodeling must be closely coordinated with ER activity. Accordingly, we demonstrated here that CSP exposure induces genes involved in UPR in Af293 and CEA17, supporting the hypothesis that the cell wall remodeling process is conserved at low and high doses of CSP and among the strains. One critical step in the GPI-anchors biosynthesis is the conversion of HMG-CoA to mevalonate, a central hub in ergosterol, heme A and ubiquinone biosynthesis (Buhaescu and Izzedine, 2007) and a limiting prerequisite for siderophore production (Yasmin et al., 2012). Although the contribution of the putative HMG-CoA reductase coding gene *hmg2* for mevalonate biosynthesis is still unknown (Yasmin et al., 2011), here we described that the exposure to high CSP concentration in both strains induced its expression. In *A. fumigatus*, *hmg1* overexpression increases siderophore production and cellular ergosterol content (Yasmin et al., 2012). Curiously, especially at lower CSP concentration, we detected repression of several siderophore and ergosterol biosynthetic genes, along with amino acid biosynthetic genes and multidrug resistance pump (MDR) coding genes, suggesting that a change in the plasma membrane composition and iron homeostasis could result from CSP exposure. In fact, we detected induction of two iron transporters (*cccA* and *fetD*) that are not related to siderophores, corroborating the imbalance in iron levels after CSP exposure. At higher CSP concentrations, even genes involved in ribosome biogenesis and translation were repressed, leading us to speculate that a metabolic rearrangement suppressing biosynthetic processes that consume a considerable amount of energy and precursors occur upon CSP exposure to favor cell wall related processes and possibly others. Actually, it was previously reported that under standard cell wall stress conditions, MpkA acts as a repressor for siderophore biosynthesis while it promotes the activation of stress responses, development and secondary metabolite biosynthesis (Jain et al., 2011).

BGC induction has already been reported as a result of the cell wall remodeling process (Jain et al., 2011; Valiante, 2015), oxidative stress (Reverberi et al., 2010) and iron availability (Wiemann et al., 2014), however, the accurate biological role of SMs in these processes is not fully understood. As SM production requires a considerable amount of energy and precursors, it is possible that BGCs may be induced by the high availability of precursors resulting from the fungistatic effects of CSP. Therefore, the metabolic rearrangement mentioned above would be a strategy to use the surplus components of primary metabolism in the biosynthesis of SMs (Sheridan et al., 2015). For instance, the product of BGC9, HAS, which was highly induced in Af293, acts as an iron chelator in response to increased iron availability (Wiemann et al., 2014), therefore it is possible that the HAS cluster was induced as an attempt to protect the cells against the deleterious effects of excessive iron accumulation. On the other hand, excessive HAS abolishes the CPE, most likely because several enzymes require iron as a cofactor. The fact that iron absence differentially affects the CPE in Af293 and CEA17 strains indicates that Af293 has a more fragile balance between iron consumption and excess and / or CEA17 has a more robust iron balancing mechanism. Overall, here we detected a significant difference regarding BGC induction between the Af293 and CEA17 strains and between the two drug concentrations in Af293, reinforcing the heterogeneity in SM production between them.

We and others previously demonstrated the essentiality of a functional MRC in *A. fumigatus* CPE (Satish et al., 2019; Aruanno et al., 2019, Valero et al., 2020). These studies implied the mitochondria in (i) the energy supply for the calcium influx to trigger the proper signaling pathways (Aruanno et al., 2019) and (ii) the ROS generation resulting in lipid peroxidation and altered plasma membrane composition around Fks1, which dampens its access to CSP (Satish et al., 2019). We reported here that the MRC genes are more strongly induced when both strains are treated with high CSP concentration, which agrees with the emergence of paradoxically growing hyphae. The fact that the induction of MRC genes is higher in Af293 represents another possible explanation for why the CPE is more pronounced in this strain. Despite the CSP-induced accumulation of mitochondrial ROS (Satish et al., 2019), we did not detect induction of the classical antioxidant-coding genes in our transcriptome. Instead, it revealed induction of BGCs, especially in Af293 treated with high CSP concentration, suggesting that the cells could be using these compounds as an alternative to cope with the stress. For instance, we described here the induction of genes in melanin BGC, a cluster positively regulated by MpkA and typically involved in *A. fumigatus* defense against oxidative stress (Valiante et al., 2016).

An increase in intracellular calcium is essential for CPE in *A. fumigatus* (Ries et al., 2017, Aruanno et al., 2019, de Castro et al., 2019). It was previously reported that the calcium-responsive TF CrzA binds to and regulate the expression of chitin synthase genes in this process (Ries et al., 2017). Here we showed that CrzA binds to promoter regions of hundreds of genes at standard conditions and that the binding intensity increases at several promoters upon caspofungin exposure. Not surprisingly, we described that among the CrzA targets are genes that belong to nearly all processes which we reported here as modulated by CSP, confirming the central role of this TF in CSP response. Although the *in silico* analysis did not detect significant differences between the CrzA binding motifs of Af293 and CEA17, our work highlights the heterogeneity of this TF in the two strains, as the *crzA* deletion impacted different sets of genes in Af293 and CEA17. Besides the growth defect observed in Δ*crzA*^Af293^, this mutant fails to activate several genes involved in the cell wall metabolism upon CSP exposure, which probably renders this mutant a thicker cell wall and CPE absence. Given the essentiality of *mpkA* for the CPE (Ries et al., 2017) and the negative effect of *crzA* deletion on MpkA phosphorylation upon CSP exposure, our work places MpkA and CrzA as the main players of CSP response, especially in Af293.

The *crzA* deletion also resulted in BGC induction in both strains, and SM production was more prominent in Δ*crzA*^Af293^ mutant, as demonstrated by our mass spectrometry studies. Here, we showed that CrzA binds to the promoter regions of most genes in siderophore BGC. As siderophores are involved in iron acquisition (Misslinger et al., 2020), and several SMs are responsive to changes in iron levels (Wiemann et al., 2014) we propose that CrzA (and calcium signaling) is involved in the iron homeostasis process, a role never reported before. Besides iron acquisition, the siderophores, along with gliotoxin, have an essential role in oxidative defense (Schrettl et al., 2007; Schrettl et al., 2010). Given the induction of MRC genes (and presumably greater mitochondrial activity and ROS generation) by CSP exposure (Satish et al., 2019; Aruanno et al., 2019; Valero et al., 2020), it is possible that the production of these SMs was a result of increased mitochondrial ROS production.

We tested FCSP and observed that in *A. fumigatus* the fluorescent drug initially accumulates in the interface cell wall / cell membrane but it is progressively internalized into vacuoles most likely by endocytosis, since possible small endocytic vesicles are observed. Jaber *et al*. (2020) have demonstrated that this FCSP also accumulates in *Candida albicans* vacuoles and cellular uptake was energy dependent and reduced by endocytosis inhibitors. We propose that FCSP vacuolar trafficking is an attempt of both *C. albicans* and *A. fumigatus* to remove the caspofungin-Fks1 (β-1,3-glucan synthase) complexes from the cell membrane by membrane turnover via endocytosis contributing to explain the previous observation that CSP induces *A. fumigatus* Fks1 relocation from the hyphal tips to the vacuole (Moreno-Velásquez et al., 2017). This probably would help to cope with CSP toxicity trying to replace the “damaged” CSP-Fks1 complexes by novel Fks1. However, as emphasized by Jaber and collaborators (2020), it is not clear if the presence of the drug in the vacuole accelerates the toxicity to CSP in both fungal species. Interestingly, we have not observed significant differences in wild-type and their corresponding *crzA* null mutants strongly indicating that both clinical strains are using common mechanisms of CSP internalization at the initial steps of interaction of the fungus with CSP.

Overall, our data suggest not only that CrzA signaling is more crucial for Af293 than for CEA17, but also that CSP exposure is more damaging for Af293. The combined stresses of *crzA* absence and CSP exposure are highly dependent on a delicate balance between high demand and economy of iron, which is apparently achieved in CEA17 but not in Af293. Our studies provide valuable comparative information about *A. fumigatus* strain heterogeneity that can be useful for the development of antifungal agents that potentiate the caspofungin activity.

## Materials and Methods

### Strains, media and drugs

All the *A. fumigatus* strains used in this work are described in Supplementary Table 5. Strains were grown in *Aspergillus* minimal medium (AMM, 1% w/v glucose, 50 mL/L of a 20x salt solution, 1 mL/L of 5 x trace elements, pH 6.5) or complete media (YG, 2% w/v glucose, 0.5% w/v yeast extract, 1mL/L of 5 x trace elements) (Kafer et al., 1977). Iron-depleted AMM were prepared in the same manner than AMM except that all iron-containing compounds were taken out from the trace elements. Agar was added to a final concentration of 2 % w/v to the AMM.

### CSP paradoxical effect (CPE) test

To determine *A. fumigatus* growth in presence of CSP in liquid media, 1.10^4^ spores/ml were dispensed into a 96-well F-bottom microplate containing 200µl of liquid AMM plus 0.2 or 2 µg/ml of CSP (Sigma). The plates were incubated at 37°C for 48 hours. To determine growth on solid media, a drop containing 1.10^5^ spores was inoculated at the center of a petri dish with solid AMM plus 1 or 8 µg/ml of CSP. The plates were incubated at 37°C for 5 days.

### RNA sequencing

Conidia (1.10^7^) were grown into liquid AMM for 16 h at 37°C under shaking conditions prior to addition of 0.2 or 2 µg/ml of CSP for 1 hour. Mycelia were harvested and frozen in liquid nitrogen. Total RNA was extracted using Trizol method (Invitrogen), treated with RNase-free DNase I (Fermentas) and purified using a RNAeasy Kit (Qiagen) according to manufacturer’s instructions. The RNA from each exposure was quantified using a NanoDrop and Qubit fluorometer and analyzed using an Agilent 2100 Bioanalyser system to assess the integrity of the RNA. RNA Integrity Number (RIN) was calculated; RNA sample had a RIN = 9.0 - 9.5. For library preparation, the Illumina TruSeq Stranded mRNA Sample Preparation kit was used according to manufacturer’s protocol. The libraries were sequenced (2×100bp) on a HiSeq 2500 instrument, sequencing approx. 17.3.10^6^ fragments per sample. Short reads were submitted to the NCBI’s Short Read Archive under the Bioproject SRP154134. Raw reads were aligned to Af293 reference genome using Hisat2 and the differential gene expression quantification was performed as described previously (Ries et al., 2020). Significant differentially expressed genes (DEGs) were selected using q-value <= 0.05 and log2(fold-change) >= 0.6 or log2(fold-change) <= −0.6. Gene Ontology enrichment analysis in DEGs was performed using Bioconductor package topGO (version 2.38.1). The RNA-seq data are available from NCBI SRA (sequence read archive) database under accession number SRP154134.

### ChIP-sequencing

Conidia were grown into liquid minimal media and treated with CSP as described above. Chromatin preparation was performed as described previously (Suzuki et al., 2012). For immunoprecipitation, 2µg of anti-GFP antibody (ab290, Abcam) was used. Library preparation was carried out using NEBNext® Ultra II Library Prep kit (Illumina, cat. no. E7645L) according to manufacturer’s protocol. Libraries were checked and quantified using DNA High Sensitivity Bioanalyzer assay (Agilent, cat. no. XF06BK50), mixed in equal molar ratio and sequenced using the Illumina HiSeq2500 platform at the Genomics and Single Cells Analysis Core facility at the University of Macau.

Raw sequencing reads were mapped to the *A. fumigatus* strain Af293 reference genome (AspGD version s03-m05-r06) using Bowtie2 (v2.3.5) (Langmead and Salzberg, 2012). For CrzA:GFP binding sites analysis, the peak calling was performed as described previously (Ries et al., 2020) and the peaks were individually checked using the Integrative genomics viewer (IGV) tool (Robinson et al., 2011). The peaks located up to 2000 bp from a gene, with log10(pvalue) > 10 were considered significant. For motif enrichment analysis, 200 and 500 bp sequences around the significant peaks were uploaded to MEME-ChIP (https://meme-suite.org/meme/tools/meme-chip) software. The ChIP-seq data are available from NCBI SRA database under accession number GSE179132.

### Promoter regions analysis

To compare CrzA binding motifs in *A. fumigatus* strains A1163 (CAE17) and Af293, we identified single-copy orthologous genes between the two strains using OrthoFinder, version 2.4.0 (Emms and Kelly, 2019). We then retrieved the corresponding non-coding region sequences of all single copy orthologs. We defined a gene’s non-coding region as up to 2kb of intergenic sequence upstream of the transcription start site, but not including sequence of the upstream gene (e.g., if there was another gene 1.5 kb upstream, we only retrieved the 1.5 kb of non-coding sequence). To search for occurrences of the CrzA binding motif, we used the Find Individual Motif Occurrences (FIMO) program, version 5.3.3 (Grant et al., 2011). We specifically investigated the occurrence and conservation of the CrzA binding motif in Af293 non-coding regions whose orthologs were shown to exhibit CrzA binding via ChIP-seq in the A1163 strain. To elucidate differences in non-coding regions between the two strains, we performed sequence alignments and visualized key sequence differences via MAFFT version 7.453 (Katoh and Standley, 2013).

### TEM analysis of cell wall

Conidia (1.10^7^) were inoculated in AMM and grown statically for 24 h at 37°C. Mycelia were treated with 0.125 µg/ml of CSP for 120 min and the control was left untreated. Mycelia were harvested and immediately fixed in 0.1 M sodium phosphate buffer (pH 7.4) containing 2.5% glutaraldehyde (v/v) and 2% paraformaldehyde (w/v) for 24 h at 4°C. Samples were encapsulated in agar (2% w/v) and subjected to fixation with 1% OsO4, contrasting with 1% uranyl acetate, ethanol dehydration, and a two-step infiltration process with Spurr resin (Electron Microscopy Sciences) for 16 h and 3 h at room temperature (RT). Additional infiltration was provided under vacuum at RT before embedment in Beem capsules (Electron Microscopy Sciences) and polymerization at 60°C for 72 h. Semithin (0.5-µm) survey sections were stained with toluidine blue to identify the best cell density areas. Ultrathin sections (60 nm) were prepared and stained again with uranyl acetate (1%) and lead citrate (2%). Transmission electron microscopy (TEM) images were obtained using a Philips CM-120 electron microscope at an acceleration voltage of 120 kV using a MegaView3 camera and iTEM 5.0 software (Olympus Soft Imaging Solutions GmbH). Cell wall thicknesses of 100 sections of different germlings were measured at ×23,500 magnification. Images were analyzed with the ImageJ software (Schneider et al., 2012). Statistical differences were evaluated using a Two-way analysis of variance (ANOVA) and Sidak’s post hoc test.

### Immunoblot analysis

Conidia (1.10^7^) were grown into liquid minimal media for 16 hours and treated with increasing CSP concentrations for 1 hour before being harvested. Protein extractions and Western blots were performed as described previously (de Assis et al., 2015). Phosphorylated and total MpkA were detected using anti-phospho-p44/42 MAPK antibody and anti-p44/42 MAPK antibody, respectively, as described previously (Ries et al., 2017).

### Microscopy

Conidia of each specific strain were grown on coverslips in 4 ml of minimal medium at 30°C. After 16h of incubation, the coverslips with adherent germlings were treated with 0.125µg/mL of a functional fluorescently labeled probe of caspofungin (FCSP) for 5, 15 and 30 min (Jaber et al., 2020). Staining of vacuoles was performed by the addition of 10 µM CellTracker Blue CMAC dye for 10 min at 30°C. Subsequently, the coverslips were rinsed with phosphate-buffered saline (PBS; 140 mM NaCl, 2 mM KCl, 10 mM NaHPO4, 1.8 mM KH_2_PO_4_, pH 7.4). Slides were visualized on the Observer Z1 fluorescence microscope using a 100x objective oil immersion lens. Differential interference contrast images (DIC) and fluorescent images were captured with AxioCam camera (Carl Zeiss) and processed using AxioVision software (version 4.8). In each experiment, at least 30 gemlings were counted. For FCSP the excitation wavelength was 560/20 nm and the emission wavelength was 629.5/37.5 nm. For CellTracker Blue CMAC, the excitation wavelength was 450 to 490 nm, and emission wavelength was 500 to 550 nm.

### High-resolution mass spectrometry analysis

Conidia (1.10^4^) were inoculated in 50 ml of liquid AMM and incubated for 24 hours at 37 ℃ under shaking conditions. After this period, 0, 0.2 or 2 µg/ml of CSP were added and the cultures were incubated under the same conditions for further 72 hours. The supernatants were frozen dried, and 100 mg were extracted with methanol in ultrasonic bath for 40 min. The extracts were then filtered at 0.22u PTFE and submitted to analysis. High-resolution mass spectrometry analyses were performed in a Thermo Scientific QExactive© Hybrid Quadrupole-Orbitrap Mass Spectrometer. Analyses were performed in positive mode with *m/z* range of 133-2000; capillary voltage at 3.5 kV; source temperature at 250 °C; S-lens 50 V. The stationary phase was a Thermo Scientific Accucore C18 2.6 µm (2.1 mm x 100 mm) column. The mobile phase was 0.1% formic acid (A) and acetonitrile (B). Eluent profile (A/B %): 95/5 up to 2/98 within 10 min, maintaining 2/98 for 10 min and down to 95/5 within 1.2 min held for 8.8 min. Total run time was 30 min for each run and flow rate 0.3 mL.min^-1^. Injection volume was 5 µL. MS/MS was performed using normalized collision energy (NCE) of 30 eV; maximum 5 precursors per cycle were selected. MS and MS/MS data were processed with Xcalibur software (version 3.0.63) developed by Thermo Fisher Scientific.

### Library search in the GNPS database and metabolite identification

**I**n order to identify secondary metabolites in *A. fumigatus* extracts using GNPS database, a high throughput dereplication of the MS/MS analyses was performed using the online workflow (https://ccms-ucsd.github.io/GNPSDocumentation/) on the GNPS website (http://gnps.ucsd.edu). The data was filtered by removing all MS/MS fragment ions within +/- 17 Da of the precursor m/z. MS/MS spectra were window filtered by choosing only the top 6 fragment ions in the +/- 50 Da window throughout the spectrum. The precursor ion mass tolerance was set to 0.02 Da and a MS/MS fragment ion tolerance of 0.02 Da. The spectra were then searched against GNPS’ spectral libraries. The library spectra were filtered in the same manner as the input data. All matches kept between network spectra and library spectra were required to have a score above 0.5 and at least 5 matched peaks (Wang et al., 2016). The results obtained in GNPS database are available at https://gnps.ucsd.edu/ProteoSAFe/status.jsp?task=3d430a0319bc414dbfd674688234e066. Yet, metabolites were also identified through MS/MS spectra previous reported in the literature.

## Acknowledgement

We thank Dr. Nancy P. Keller for providing the secondary metabolite mutants.

---

**Supplementary Figure 1:** Scatter plots representing the correlation between the relative expression of the transcriptome pairs.

**Supplementary Figure 2:** Radial growth in solid media of Wt, Δ*fapR*, Δ*fmaA* and Δ*hasA* muntants exposed to 1 and 8 µg/ml of CSP.

**Supplementary Figure 3:** Cellular localization determined by microscopy of FCSP in the Af293, Δ*crzA*^Af293^ and Δ*crzA*^CEA17^ strains after 5, 10 and 15 minutes of exposure to 0.125 µg/ml, the vacuoles were labeled with blue CMAC dye.

**Supplementary Figure 4:** Relative expression of genes involved in cell-wall metabolism in Af293, CEA17, Δ*crzA*^Af293^ and Δ*crzA*^CEA17^ exposed to LD and HD of CSP.

**Supplementary Table 1:** Relative expression of selected genes from Af293 and CEA17 after exposure to LD and HD of CSP.

**Supplementary Table 2:** Relative expression of 26 BGCs from Af293, CEA17, Δ*crzA*^Af293^ and Δ*crzA*^CEA17^ strains exposed or not to LD and HD of CSP.

**Supplementary Table 3:** CrzA targets in CEA17 before and after exposure to HD of CSP.

**Supplementary Table 4:** Go terms enriched among the genes induced and repressed in Δ*crzA*^Af293^ and Δ*crzA*^CEA17^.

**Supplementary Table 5:** Strains used in this work

**Supplementary File 1:** Relative expression (log_2_ fold-change) of all *A. fumigatus* genes in each transcriptome.

